# Sex, males, and hermaphrodites in the scale insect *Icerya purchasi*

**DOI:** 10.1101/2020.09.04.281618

**Authors:** Andrew J. Mongue, Sozos Michaelides, Oliver Coombe, Alejandro Tena, Dong-Soon Kim, Benjamin B. Normark, Andy Gardner, Mark S. Hoddle, Laura Ross

## Abstract

Androdioecy (the coexistence of males and hermaphrodites) is a rare mating system for which the evolutionary dynamics are poorly understood. Here we study the only presumed case of androdioecy in insects, found in the cottony cushion scale, *Icerya purchasi*. In this species, female-like hermaphrodites have been shown to produce sperm and self-fertilize. However, rare males are sometimes observed too. In a large population-genetic analysis, we show for the first time that although self-fertilization appears to be the primary mode of reproduction, rare outbreeding events between males and hermaphrodites do occur, and we thereby confirm androdioecy as the mating system of *I. purchasi*. Thus, this insect appears to have the colonization advantages of a selfing organism while also benefitting from periodic reintroduction of genetic variation through outbreeding with males.

## Introduction

Reproductive systems are remarkably variable across life, yet this variation is poorly understood in evolutionary terms. The forces that favor different modes of reproduction, as well as those that shape taxonomic and phylogenetic distributions remain elusive. The evolutionary and zoological literature has predominantly focused on the distinction between sexual and asexual reproduction (Innes et al. 2000; Silvertown 2008; Gibson et al. 2017). However, much of the variation in reproductive systems is found among those that reproduce sexually (White 1973; Charnov et al. 1976; Bachtrog et al. 2014). For example, although most animals are gonochoristic (i.e. have separate sexes) with genetic sex determination, more than 20% of species have a different reproductive system (Tree of Sex Consortium 2014). The evolution of many of these reproductive systems remains poorly understood.

One such system is androdioecy, which is characterized by the coexistence of males and hermaphrodites and the absence of females. Androdioecy is a rare mating system. To date, it has been found in only 115 species of animals (Weeks 2012), including just three species of vertebrate, all fish (Smith 1965; Taylor et al. 2001), and a small number of invertebrates (e.g. the nematode *C. elegans*). Outside of the animal kingdom, it has also been found in several plant species (Lloyd 1975; Liston et al. 1990; Pannell 1997). As such, androdioecy appears to have independently evolved several times in eukaryotes and is often thought of as an advantageous strategy for colonizing new locations: having the benefit of reproductive assurance (i.e. allowing even a single hermaphrodite to found a new population through self-fertilization) without the inbreeding cost of pure selfing as the production of males permits outcrossing (Weeks 2012). Yet this logical hypothesis has not been formally tested and it remains unclear how organisms anatomically and cytologically acquire the capacity to produce both gamete types in one phenotype but not the other.

In plants, androdioecy is considered an intermediate condition in the evolutionary transition from hermaphroditism to separate sexes. In animals however, most transitions appear to go in the opposite direction (from separate sexes to hermaphroditism then androdioecy, e.g., barnacles Høeg 1995). Another difference between plant and animal androdioecy is that, in animals, hermaphrodites are typically unable to mate with each other but are able to either self-fertilize or mate with males (Weeks et al. 2006). This suggests either that the evolutionary forces shaping the evolution of androdioecy are different in the two kingdoms or that the evolution of outbreeding hermaphrodites is constrained differently by the anatomy of plants and animals. Indeed, it has been argued that the evolution of cross-fertile hermaphroditic animals poses a greater evolutionary challenge as it would require not only changes to gamete production, but also novel rearrangement of the reproductive tract to allow for bidirectional copulation (Pannell 2002; Weeks et al. 2006, 2009). Moreover, plants often possess multiple, independently formed reproductive structures (i.e. flowers), compared to the single-origin reproductive structures of animals. It stands to reason that with the evolution of novel reproductive strategies, plants may rescue some fitness from sub-functional reproductive organs, while in animals, sterility tends to be a binary trait. As such, androdioecy in animals may be the more tractable evolutionary route to outcrossing and the presence and frequency of males can be understood in terms of selection for outbreeding opportunity. However, to assess these hypotheses requires first identifying and characterizing potential cases of androdioecy in the animal kingdom for further study.

Insects are the most species-rich clade of animals and display exceptionally variable reproduction, including chromosomal sex determination, haplodiploidy, and – very rarely – hermaphroditism (de la Filia et al. 2015; Blackmon et al. 2017). The only known cases of hermaphroditism in insects are found in a tribe of scale insects (plant-feeding insects in the order Hemiptera), the Iceryni. Hermaphroditism has been recorded in three species within this tribe, each of which appear to constitute an independent origin of hermaphroditism (Unruh and Gullan 2008; Tree of Sex Consortium 2014). Hermaphroditic individuals display a female phenotype (scale insects display strong sexual dimorphism with relatively large wingless females and very small winged males, see figure 1a&b) and appear incapable of mating with each other, despite containing an ovitestis that produces both sperm and eggs. This suite of phenotypes matches the expectation for a hermaphroditic animal described above, but there is some evidence that these species are actually androdiecious. First of all, true males are observed in natural populations (the male in figure 1b is from one of these hermaphroditic species, *Icerya purchasi*), although they are rare (Hamon and Fasulo 2005; Kim et al. 2011), with reported frequencies differing substantially between populations (from 0.01%-10%). Secondly, these rare males can mate with the hermaphrodites (figure 1c). However, if males do not contribute genetically to the next generation, their presence is irrelevant from a reproductive perspective and the system is, genetically at least, purely hermaphroditic. It has been unclear whether these rare males are indeed able to fertilize hermaphrodites, whether their sperm is able to compete with the sperm produced by the hermaphrodite and, if this is the case, how important outbreeding events are in natural populations.

**Figure 1.**
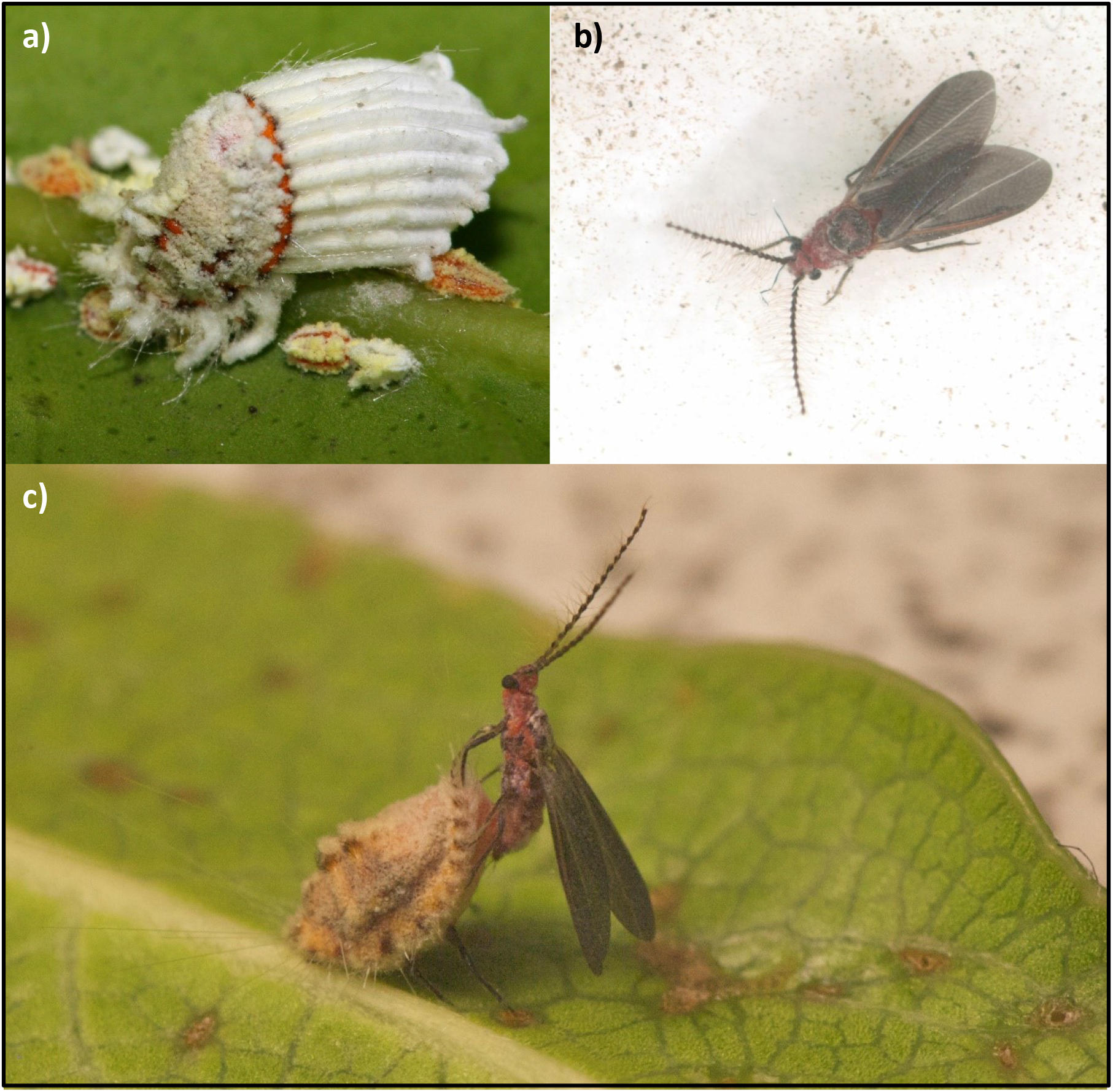
*Icerya purchasi* hermaphrodites (with white egg case and crawlers) are most commonly observed **(a)** but rare males have been reported **(b)** and we have observed matings between the two sexes under laboratory conditions, with the hermaphrodite enabling mating by raising their body (image credit to Enric Frago) **(c)**. It is presently unclear if these matings lead to the fertilization of eggs.

The presence of androdioecy in Iceryini would be remarkable as it would not simply be the only known instance of this mating system among insects, but also the first record of androdioecy having evolved from haplodiploidy (in which females develop from fertilized eggs and males from unfertilized eggs (Schrader and Hughes-Schrader 1926)). Cytogenetic studies have shown that while hermaphrodites are diploid (2n=4), both males and the sperm-producing section of the ovitestis within hermaphrodites are haploid (n=2, Kokilamani et al. 2019). It has been suggested that the evolution of androdioecy in the Iceryini could result from conflict between males and females over the proportion of eggs that are fertilized, a conflict that only exists in haplodiploid species (Normark 2009; Gardner and Ross 2011).

The presence of an unusual haploid tissue within the ovitestis of a female-like hermaphrodite is a unique feature of Iceryini that is poorly understood. The first study that described this phenomenon suggested that the haploid tissue arises due to the (random) loss of one haploid copy of the genome (Hughes-Schrader 1925) from germline cells destined to produce sperm (Figure 2a). However, a later study suggested that instead this haploid tissue originates from supernumerary sperm cells transmitted by males during mating (Royer 1975). Cytogenetic analyses show that oocytes are penetrated by multiple sperm cells, one of which fertilizes the eggs and gives rise to the diploid hermaphrodite while the other sperm cells divide to give rise to a haploid male germline (Figure 2b). By “infecting” females with sperm cells that form male gametes inside the female offspring, males effectively mate not only with the female, but also with her daughters, ensuring reproductive fitness for generations to come. A kin selection analysis of this hypothesis has suggested that this could lead to the evolution of androdioecy, although this model – which assumes no inbreeding depression or other disadvantage to offspring produced by selfing – also predicts the complete disappearance of males and cessation of outbreeding events (Gardner and Ross 2011).

**Figure 2.**
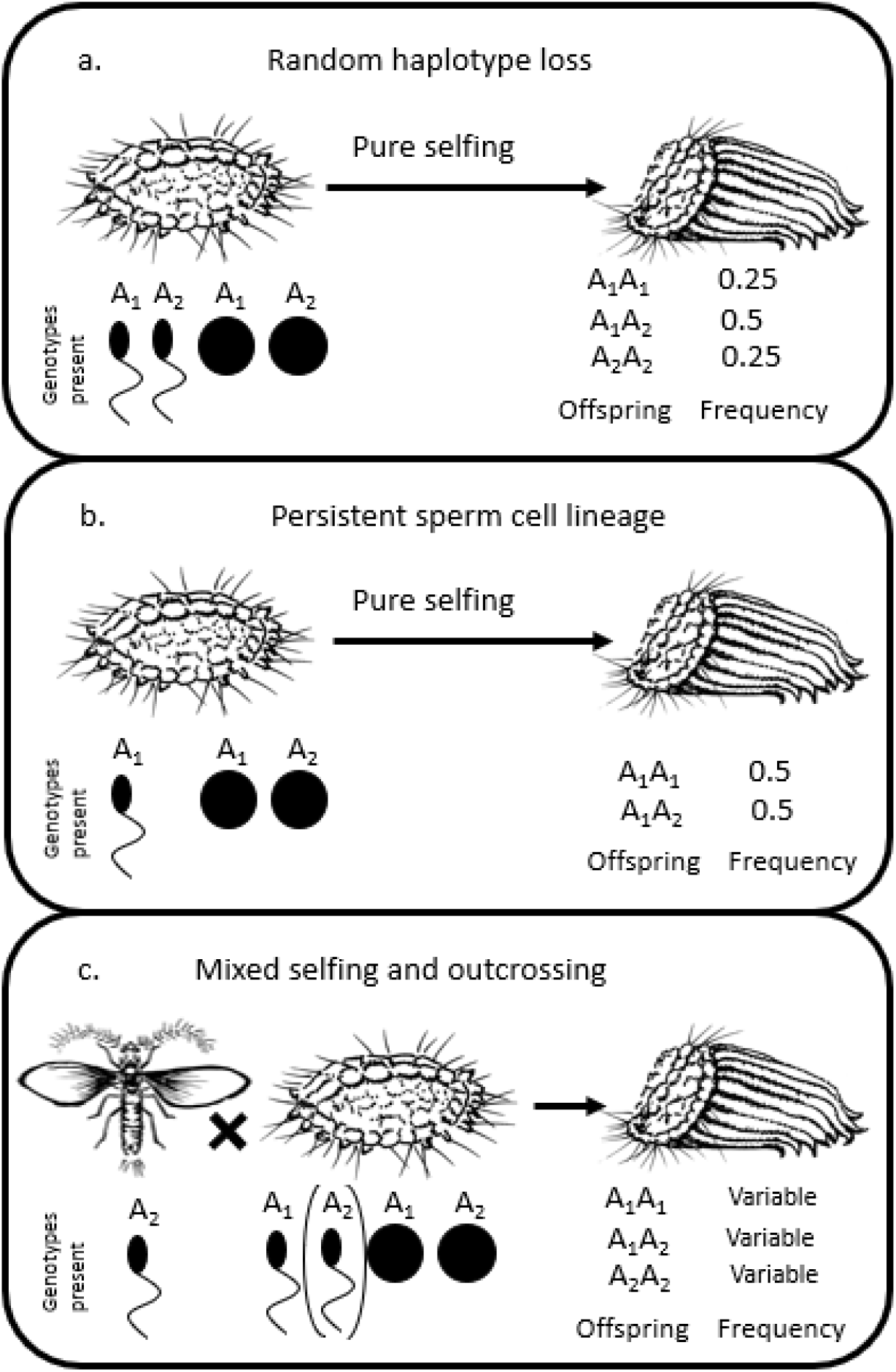
Hypothesized mating systems for *Icerya purchasi*. **a.** Sperm are formed within the diploid hermaphrodite by traditional meiotic processes resulting in all possible genotypes at Mendelian ratios. **b.** A historic outbreeding event with a haploid male results in the fertilization of an egg nucleus plus the establishment of a sperm cell lineage that persists over generations. Only two of three allelic combinations are possible under this scenario. **c.** Rare males mate with hermaphrodites, fertilizing some to proportion of their eggs, resulting in offspring of all possible genotypes with frequencies depending on the fertilization success rate of males. Note that in this scenario either selfing mechanism (a or b) would give equivalent offspring results.

These two possible mechanisms of haploid male germ-tissue formation in hermaphrodites should be easily distinguishable from each other in a purely selfing system (see expected offspring genotypes in Figure 2a&b). However, if males genetically contribute to the next generation, the inference is more complicated (Figure 2c). Thus, we first need a better understanding of the mating system of Iceryini and the role of males in reproduction. Here we focus on one of the three species of hermaphroditic scale insects in the Iceryini: the cushiony cotton scale insect, *Icerya purchasi*. This species is a highly polyphagous pest, feeding on urban ornamental plants (e.g. *Magnolia*), agricultural crops such as *Ficus* and *Citrus* (Singh and Kaur 2015; Liu and Shi 2020), and plants of conservation importance (e.g. Galapagos endemics threatened by the introduction of *Icerya*, Hoddle et al. 2013). *I. purchasi* spread from its native range in Australia over a century ago and it is now globally distributed. Climate models predict changing environmental conditions will permit further geographic expansion in the future (Liu and Shi 2020).

Given its unusual biology as well as its economic and environmental importance as an invasive pest species, insights into the population genetic structure and reproductive system of *I. purchasi* will be of interest both from an evolutionary and an applied perspective. To investigate this, we developed a panel of polymorphic microsatellite markers for use in a population genetic analysis to test for the presence of androdioecy and estimate outbreeding rates in natural populations of *I. purchasi*. Hermaphrodites are able to self-fertilize and therefore most of the population should consist of highly homozygous strains. However, if the mating system is androdiecious, we would also expect to find some heterozygous individuals resulting from recent outbreeding events. We therefore estimated outbreeding rates by examining the difference between allele and genotype frequencies in multiple populations collected globally. Finally, we used genotype data of families of wild hermaphrodites and their eggs to infer the most likely mechanism of male-germline formation in hermaphrodites (Figure 2).

## Methods

### Specimen collection

We initially collected 343 adult and 536 egg samples from 27 locations in 11 countries on 5 continents (see table S2 and figure S1). When possible, we collected hermaphrodites as well as their egg masses and these specimens were stored in individual tubes in 99% ethanol at −20°C. We were unable to obtain male specimens from field populations, either because they are rare, or because the short lifespan of adult males makes them hard to detect under field conditions, or both. For each hermaphrodite we dissected the tip of the head (<1mm) to use for the analysis. This avoided the inclusion of the ovitestis and allowed us to distinguish between a true heterozygous female or a homozygous female fertilized by and carrying eggs from a male with a different genotype. For each mature hermaphrodite that had an egg mass (white waxy ovisac, see Figure 1a), we also analyzed between one and ten embryos. These embryos were obtained by opening the egg mass attached to the hermaphrodite.

### Microsatellite markers

For the purpose of this study we designed a set of 12 polymorphic microsatellite markers. We identified the markers based on one lane of 454 (titanium) sequencing data from a pooled sample of a hermaphrodite that came from a laboratory population in California. This resulted in 94,553 reads that could be assembled to 1,772 contigs with an average length of 909 base pairs. We used the software package msatfinder (Thurston and Field 2005) to detect microsatellite repeats in the sequence data. We selected only those primers that had a tri-nucleotide repeat and had more than 10 repeats. This resulted in 40 possible microsatellite loci. We tested each of these primers under standardized PCR conditions with an annealing temperature of 55 °C. Those primers that amplified where tested for polymorphism on 6 individuals (3 from a UK and 3 from a French population). Based on these tests, we selected 12 polymorphic primers (details of these primers can be found in table S1). In order to speed up the analysis we also designed a multiplex method so that the 12 primers could be analyzed in 4 PCR 10 μL reactions and 3 ABI runs (see table S1).

### Microsatellite analysis

DNA was extracted by using the Qiagen DNAeasy kit (Hilden, Germany). Microsatellite loci were amplified with the primers described in Table S1 and repeat length was assessed via the microsatellite plugin in Geneious R8 (Kearse et al. 2012). We only counted alleles that were found in at least two separate individuals. The embryos we analyzed were very small and so it was not always possible to get enough DNA for a reliable analysis. In order to avoid calling erroneous alleles in low quality samples we removed samples in which fewer than two thirds of the loci amplified.

### Data analysis

We carried out two sets of analyses based on microsatellite genotyping of *I. purchasi*. First, we considered wild caught hermaphrodites from populations around the world and estimated allele frequencies, using the R package *polysat* (Clark and Jasieniuk 2011) to calculate Fst and create an input for population structure analyses. For this objective, we considered only individuals with full genotype scores at each locus (n = 208 across 8 countries) and generated 10 independent runs of population structure inference from k = 2 to 10 populations (Pritchard et al. 2000). We aggregated runs and used the webtool CLUMPAK to infer the best k (Kopelman et al. 2015). For F_ST_ estimates, we tested for an effect of isolation by distance, using a simple linear regression between pairwise population differentiation and physical distance between the two populations. We used shortest distances (i.e. straight lines, over water if necessary) to when comparing two locations. For countries with multiple sampling locations, we averaged the latitudes and longitudes of sampling locations within the country. To estimate per-population selfing rate (via the inbreeding rate, F), we used the R package *adegenet* (Jombart 2008) on individuals with at least 2/3rds of loci amplifying ( n = 297 across 10 countries); this function implements a simple Mantel test to calculate identity by descent. All analyses were carried out in R version 3.6.0 (Team 2019).

Second, we considered the subset of adult hermaphrodites from which we also collected and genotyped eggs. In these analyses, we compared parent and offspring genotypes to look for evidence of recent outbreeding. More specifically, we examined cases in which offspring had either: alleles not present in their sequenced parent and/or genotypic combinations not present in the parent. For these analyses, we included only parent-offspring sets in which at least half of the loci were genotyped.

Owing to the low local and global genetic variation, we did not observe any offspring with unique alleles absent in their hermaphroditic parent. Therefore, we focused on assessing the kinds and frequencies of genotypes of the offspring. For these analyses, we limited our dataset to families with 8 or more eggs. We counted observed genotype frequencies in offspring and used a simple X^2^ goodness-of-fit test on each locus to demonstrate the deviation from expected genotype frequencies under pure selfing with Mendelian segregation of alleles. We present the proportion of loci deviating from this expectation as further evidence for outbreeding between hermaphrodites and males. This analysis relies on the dubious assumption that all sampled loci are unlinked, but without a genome assembly to place our markers we do not have a means of excluding linked loci.

## Results

### Population genetic variation

We analyzed 208 individuals from 8 countries. On average we found 2.67 alleles per locus across 12 microsatellites, indicating low levels of variation. For instance, the samples from Korea did not show any genetic variation, all sharing a single haplotype. Likewise globally, one haplotype dominated our samples with 46% of all specimens analyzed having the same genotype across all 12 markers. The next most common haplotype, which differs at a single locus from the first, accounts for another 44% of the samples. Consequently, 90% of our sampled individuals were identical across 11 of 12 microsatellite loci.

Population structure analyses indicated the existence of 3 distinct populations, based on the biggest change in likelihood between sequential k values. However, these three clusters simply reflect the pattern described above, with one inferred population for each of the two dominant haplotypes and a third for the rarer haplotypes (Figure S2). The relationships between populations can be seen more easily in Table S3, in which we have calculated F_ST_ across each pair of sampled countries for which we have full genotyping information. Genetic differentiation is low but variable, and ranges from 0.001 to 0.1118, with no signal for isolation by distance (r^2^ = 0.002, p = 0.84).

### Outbreeding rate

Evidence for outbreeding was assessed in two ways; first, we used population genetic packages in R to estimate per-population inbreeding rates (Figure 3) based on differences between observed and expected heterozygosity. These analyses suggest that selfing is pervasive, but not complete. The global mean F is 0.642 and a majority of individuals have F > 0.5. However, a minority of individuals show quite low F-values (< 0.2), as expected if rare males create occasional outbreeding events.

**Figure 3.**
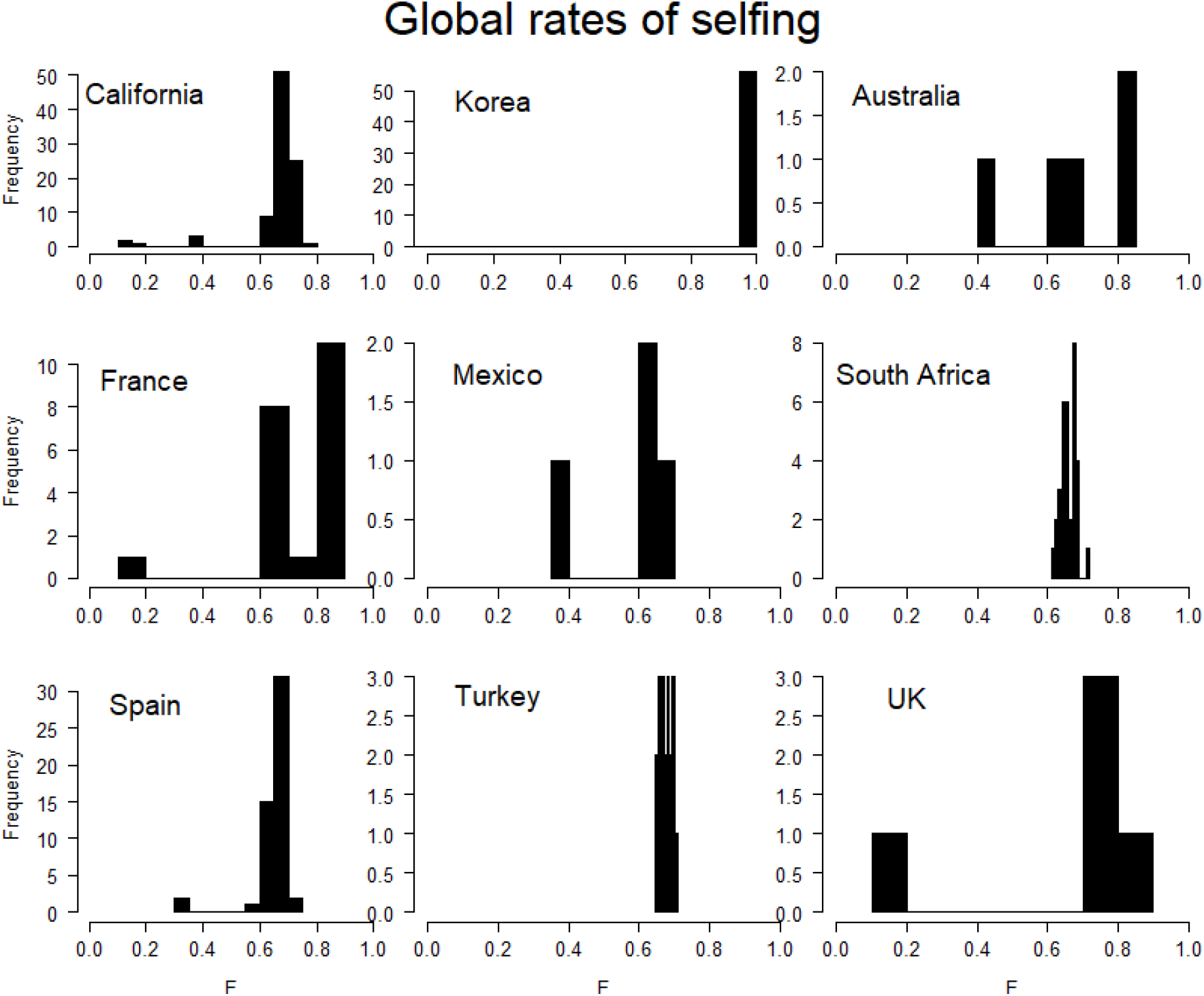
Inferred inbreeding coefficient (F) for individuals across 9 populations. Here we included only individuals for which at least two thirds of loci (8 of 12) amplified. Populations each show a mean F above 0.5 but many (e.g. Spain and France) also contain a minority of individuals showing relatively little evidence for inbreeding. Note that owing to a lack of clear population structuring, we pooled sampling locations within countries (namely, California in the United States, France, and Turkey).

Second, we examined the alleles in known parent-offspring sets. Of 536 eggs genotyped here, 96 had at least one difference from the parent’s genotype. We examined these parent-offspring sets for genotypic combinations that could distinguish the two possible mechanisms of selfing (Figure 2): Under the persistent-sperm-lineage hypothesis a selfing hermaphrodite can transmit either of two possible alleles (A1, A2) through their eggs, but only one through their sperm (A2). Thus, half of the offspring will be heterozygotes (A1, A2), and half will be homozygous for the sperm genotype (A2A2), but never the alternate homozygote (A1A1). However under the genome loss hypothesis, assuming that loss is random, a selfing hermaphrodite can transmit either of two possible alleles (A1, A2) through both their sperm and their eggs, so half the offspring will be heterozygotes (A1, A2), a quarter will be homozygous for the sperm genotype (A2A2) and remainder will be homozygous for (A1A1). We started by examining families with a heterozygous parent that showed offspring with all three genotypic combinations. Unfortunately, many of these families were represented by fewer than five eggs, which hindered our ability to estimate genotype frequencies. Of the three families with > 8 offspring sampled, two failed to amplify at multiple loci. Here we present a single family from France (one hermaphroditic parent and 10 eggs, Table 1) with the most complete genotyping. As mentioned above, expectations from selfing with a persistent sperm cell line are that only one of the two homozygous states should be found in the offspring. Thus, this system alone cannot explain the observed alleles.

**Table 1.** Investigation of offspring genotypes from a wild-caught hermaphrodite from France. Rows represent the diploid genotype (in terms of microsatellite repeat length) for an individual at each polymorphic locus (non-variable loci omitted), with observed genotype frequencies for each locus at the bottom. Under pure selfing with a persistent sperm lineage, only one of two homozygous genotypes should be observed. Likewise, under pure selfing with random haplotype loss in spermatogenesis, genotypic ratios should follow Mendelian expectations (0.25: 0.5: 0.25). Eight of nine loci have genotype frequencies (in bold and underlined) that significantly differ from this expectation under a X^2^ goodness-of-fit test. Thus, neither of these pure selfing scenarios matches observed offspring genotypes, suggesting a role for paternity from an external male for some offspring.

| Locus: | Plcp21 |  | Plcp39 |  | Plcp45 |  | Plcp71 |  | Plcp58 |  | Plcp62 |  | Plcp101 |  | Plcp37 |  | Plcp93 |  |
| --- | --- | --- | --- | --- | --- | --- | --- | --- | --- | --- | --- | --- | --- | --- | --- | --- | --- | --- |
| A<sub>1</sub> length |  |  |  |  |  |  |  |  |  |  |  |  |  |  |  |  |  |  |
| A<sub>2</sub> length |  |  |  |  |  |  |  |  |  |  |  |  |  |  |  |  |  |  |
| Parent | 183 | 186 | 171 | 165 | 173 | 179 | 99 | 108 | 131 | 137 | 173 | 176 | 98 | 107 | 206 | 215 | 115 | 121 |
| Egg1 | 183 | 186 | 165 | 165 | 179 | 179 | 99 | 108 | 137 | 137 | 173 | 173 | 98 | 98 | 206 | 206 | 115 | 115 |
| Egg2 | 183 | 186 | 171 | 171 | 173 | 179 | 99 | 108 | 131 | 131 | 176 | 176 | 98 | 107 | 215 | 215 | 121 | 121 |
| Egg3 | 183 | 183 | 171 | 165 | 173 | 179 | 99 | 99 | 131 | 137 | 173 | 176 | 98 | 107 | 206 | 215 | 115 | 121 |
| Egg4 | 183 | 186 | 165 | 165 | 179 | 179 | 99 | 108 | 137 | 137 | 173 | 173 | 98 | 107 | 206 | 206 | 115 | 115 |
| Egg5 | 183 | 183 | 165 | 165 | 179 | 179 | 99 | 99 | 137 | 137 | 173 | 173 | 98 | 98 | 206 | 206 | 115 | 115 |
| Egg6 | 183 | 183 | 165 | 165 | 179 | 179 | 99 | 99 | 137 | 137 | 173 | 173 | 98 | 107 | 206 | 206 | 115 | 115 |
| Egg7 | 183 | 186 | 165 | 165 | 179 | 179 | 99 | 99 | 131 | 137 | 173 | 173 | 98 | 98 | 206 | 206 | 115 | 115 |
| Egg8 | NA | NA | NA | NA | NA | NA | 99 | 108 | 131 | 137 | 173 | 173 | 98 | 98 | 206 | 206 | 115 | 115 |
| Egg9 | 183 | 183 | 165 | 165 | 179 | 179 | 99 | 99 | 131 | 137 | 173 | 173 | 98 | 98 | 206 | 206 | 115 | 115 |
| Egg10 | 183 | 186 | 171 | 165 | 173 | 179 | 99 | 108 | NA | NA | 173 | 176 | 98 | 107 | 206 | 215 | 115 | 121 |
| A<sub>1</sub> A<sub>1</sub> | **<u>0.44</u>** |  | **<u>0.11</u>** |  | **<u>0.0</u>** |  | **<u>0.5</u>** |  | 0.11 |  | **<u>0.7</u>** |  | **<u>0.5</u>** |  | **<u>0.7</u>** |  | **<u>0.7</u>** |  |
| A<sub>1</sub> A<sub>2</sub> | **<u>0.56</u>** |  | **<u>0.22</u>** |  | **<u>0.33</u>** |  | **<u>0.5</u>** |  | 0.44 |  | **<u>0.2</u>** |  | **<u>0.5</u>** |  | **<u>0.2</u>** |  | **<u>0.2</u>** |  |
| A<sub>2</sub> A<sub>2</sub> | **<u>0.0</u>** |  | **<u>0.67</u>** |  | **<u>0.67</u>** |  | **<u>0.0</u>** |  | 0.44 |  | **<u>0.1</u>** |  | **<u>0.0</u>** |  | **<u>0.1</u>** |  | **<u>0.1</u>** |  |

We calculated genotypic frequencies at each polymorphic locus. Under the random haplotype elimination hypothesis, selfing occurs through independent meiosis of egg and sperm cells within the hermaphrodite, generating both possible haplotypes in both sperm and egg, and yielding Mendelian genotype ratios. Expectations significantly differ from these ratios at 8 of 9 polymorphic loci (Table 1). Thus, it is likely that some portion of the offspring are the result of the external introduction of one of the alleles from mating with a male.

## Discussion

Three species of scale insects in the tribe Iceryini, *Icerya purchasi* (Royer 1975), *Auloicerya acaciae* (Gullan 1986), and *Crypticerya zeteki* (Normark 2003), are hypothesized to be largely hermaphroditic. However, the precise inheritance system for each species remains unclear. Here we examine the reproductive system of *I. purchasi* in detail. Under pure selfing, hermaphrodites can self-fertilize their eggs with their own sperm for an indefinite number of generations, rapidly exhausting genetic variation and leading to high levels of homozygosity and predictable genotype frequencies in offspring. However, the observations of occasional males raises the possibility that some portion of offspring are the result of outbreeding events between hermaphrodites and these males. To investigate this possibility, we carried out a series of population genetic analyses with microsatellite markers on wild caught *I. purchasi*.

The inbreeding coefficient (F) provides an effective estimate of the rate of successful hermaphrodite x male matings in the wild. From these estimates, we see that inbreeding (or more appropriately, selfing) rate is substantial (> 0.6) but not complete. Moreover, there is considerable variation in the rates within populations, as would be expected if some sampled individuals are derived from more recent outbreeding events than others. However, this pattern could also arise from completely selfing populations if some other mechanism maintains heterozygosity.

First, it is worth considering that, without a better understanding of the genomic features of *I. purchasi*, it is possible that at least some of our markers are subject to selective rather than neutral forces. If some of our microsatellites are linked to recessive lethal variants, then we might expect variation to be maintained at low levels via selection against homozygotes at these loci (Kimura et al. 1963). However, we have no reason to think our markers should be effectively non-neutral, and moreover, given the substantial amount of selfing, homozygous genotypes should frequently arise and genetic load should be effectively purged by selection in a few generations (Fox et al. 2008), as has been seen with inbreeding in other insects (Haikola et al. 2001; Mongue et al. 2016). Second, consider that the observed heterozygosity may not be due to the maintenance of alleles, but *de novo* mutation. Microsatellite repeats are known to be more variable than point mutations, with mutation rate estimates as high as 10^-3^ per gamete per generation (Brinkmann et al. 1998; Schlötterer et al. 1998). But even this high rate of mutation would not be enough to generate the number of heterozygous loci we observe in a small sample of *I. purchasi*. Moreover, there is some evidence that microsatellites generally evolve faster in species with monocentric chromosomes (Jonika et al. 2020), and mutation rate is positively related to genome-wide heterozygosity (Amos 2016). Given that *I. purchasi* is a holocentric species (Melters et al. 2012) with very low heterozygosity, the expectation is that microsatellites should evolve slower than average. Thus, although our observations are consistent with either of the above scenarios, they both strain credulity.

A much more plausible explanation is the reintroduction of heterozygosity into lineages via outcrossing between hermaphrodites and males, in other words, androdioecy.

Further suggestion of outbreeding comes from the examination of a wild-caught hermaphrodite and ten of its eggs. Considering heterozygous loci in the parent, the presence of all three genotypic combinations in the offspring rules out a system of pure selfing with a persistent sperm lineage, which would generate only two of the three genotypes. Next, considering genotype frequencies at heterozygous loci, we would expect a Mendelian ratio of homozygotes and heterozygotes if sperm are created by the random meiotic elimination of one of the two alleles during spermatogenesis. On the contrary, all nine heterozygous loci deviate from this expected ratio, eight of them significantly so. Thus, these offspring do not appear to be the result of this model pure selfing either. Rather, the skewed genotype frequencies observed can be explained by a mix of one of the two selfing systems above and fertilization from an outside male, which is androdioecy.

### *Population structure of* Icerya purchasi

As predicted for a species with a substantial selfing-rate, *I. purchasi* populations are highly homozygous, with 90% of samples having an identical diploid genotype at 11 of 12 loci. Structure analyses are hindered by this homogeneity, but our analyses of worldwide genetic variation in *I. purchasi* corroborate the narrative of global expansion through repeated, (presumably) accidental introductions by humans into new areas. No signal of isolation by distance is detectable. For example, South Africa and South Korea have nearly identical genotypes, (F_ST_ = 0.0001), whereas South Africa and France (F_ST_ = 0.1118) have the highest divergence of any pair of populations. And more generally, genetic differentiation does not vary predictably with physical distance. Presence of identical haplotypes around the world might suggest that these populations share a common origin (i.e. derived populations all arise from the same ancestral population rather than secondary spread from introduced populations), but a much larger sample of loci (e.g. from whole-genome SNP data) will be needed to reconstruct within-species relationships of the global populations to confirm or refute this possibility.

Finally, *I. purchasi* is thought to be Australian in origin (Prasad 1989) and, consequently, one might expect genetic diversity to be highest there. Speaking to this, there are at least ten private alleles not seen in any other population from this initial sample. Needless to say, it is difficult to accurately assess this pattern with a mere twelve loci from seven individuals. As above, a larger set of genetic markers and increased sampling across this native range will be needed to fully resolve the population genetics.

### Androdioecy in ecological context

Although the present genetic markers do not permit us to resolve the colonization history of this insect, it is clearly globally distributed and androdioecy likely contributed to its colonizing success following accidental introduction into new areas by humans (most likely through trade in live plants). Its ability to self-fertilize allows a single individual to establish a new colony on its own, while occasional outbreeding can maintain variation on which selection can act, allowing it to adapt to new ecological conditions faster than clonal reproduction would allow.

This system, although physiologically distinct from that of (fellow hemipteran) aphids, bears striking ecological similarity to their parthenogenesis with occasional sexual reproduction (Blackman 1980). In each, a single individual has the ability to found a new population: either through selfing in *I. purchasi* or through the parthenogenetic production of daughters in aphids. Likewise, both systems retain a capacity for sexual reproduction via mating with rare males. Scale insects and aphids share a similar phytophagous lifestyle, so it is worth considering that selective pressures to exploit the geographic and phylogenetic range of host plants have produced complimentary reproductive dynamics in these herbivores. Understanding these systems is thus important not only for evolutionary theory, but also for the development of management strategies for these crop pests.

### Conclusions

Here we have presented evidence, both at the population and pedigree scale, that *I. purchasi* is the first confirmed androdiecious insect. Self-fertilization of hermaphrodites appears to be the most common mode of reproduction, leading to low levels of genetic variation at (putatively) neutral markers. Yet both physical observation and genetic data reveal that males occasionally mate with and fertilize hermaphrodites, thus preventing a complete loss of genetic variation. This mixed system defies clear predictions for the presence and frequencies of genotypes in offspring, thus it remains an open question with what cytogenetic mechanism hermaphrodites self-fertilize. Nevertheless, this corroborating evidence of androdioecy lays the groundwork to investigate how males are produced and why they vary substantially in frequency between populations, how this rare and unusual system of reproduction evolved, and implications for managing this destructive invasive pest.

## Supporting information

Supplementary results

## Acknowledgements

The authors would like to thank David Haig and Stuart West for initial discussions, Lorenzo Santorelli, Enric Frago, and Aniek Ivens for help and advice in the lab, and members of the Ross lab for further discussion and comments. In addition, the authors thank Alicia Woods for creating and permitting use of the drawings of *Icerya purchasi* in Figure 2 and Enric Frago for the photograph in Figure 1c. This work was supported by a number of fellowships, namely a University Research Fellowship from Royal Society of London and a Junior Research Fellowship from Balliol College, Oxford. Funding came from Independent Research Fellowships from Natural Environment Research Council (grant no. NE/K009524/1 to AG and NE/K009516/1 to LR), a Consolidator Grant from European Research Council (grant no. 771387 to AG), a European Research Countil Starting Grant (PGErepro, to LR) and a Royal Society Newton fellowship (to LR).

