## Supplementary results for "Sex, males, and hermaphrodites in the scale insect *Icerya purchasi*"

**Appendix: population genetic considerations in the cottony cushion scale insect**

*Inbreeding coefficients under the Icerya purchasi mating system*

Methods developed to estimate the effect of self-fertilization on the inbreeding coefficient of an individual $F_I$ are based on individuals with diploid male and female germline. Take for example an outbred individual with a genotype of A1A2 that reproduces by selfing. The chance that its offspring is homozygous (for alleles identical by decent) is the chance that it is either A1A1 (1/4) or A2A2 (1/4) which is ½. When these offspring in turn self-fertilize the inbreeding coefficient of their offspring is a $½+½F_{t-1}$, where $F_{t-1}$ is the inbreeding coefficient of its parent. In fact, under self-fertilization the rate of inbreeding is constant and the change over the generations is described by $F_t= ½ (1+ F_{t-1})$. This predicts that under selfing the inbreeding coefficient exceeds 0.95 after just 5 generations. The question is now if the same equation holds for the mating system of *Icerya purchasi*. Consider an outbred hermaphrodite with genotype A1A2 that resulted from a double fertilization event. If we assume that A2 was the paternal allele, then this individual will produce female gametes that can either be A1 or A2 while producing male gametes with the genotype A2. A fusion of its male and female gametes will therefore result in ½ A1A2 and ½ A2A2 offspring which leads to an inbreeding coefficient of ½. The next generation of inbreeding leads to genotype frequencies of ¼ A1A2 and ¾ A2A2. This shows that the inbreeding coefficient under selfing in *Icerya purchasi* follows the same equation that has been previously derived for diplodiploid selfing. This means that it is valid to use the well-developed framework for estimating selfing rate from population level estimates of inbreeding in *Icerya purchasi*.

*Local fixation of alleles*

Although inbreeding does not influence global allele frequencies it might affect the chance that an allele becomes fixed in a local population. Under pure self-fertilization in a diplodiploid individual this is generally not the case, as a local population founded by a A1A2 hermaphrodite will converge to a population that consists of a mixture of A1A1 and A2A2 individuals. This is not the case for *Icerya purchasi*, in which A1A2 individuals (with a A2 male germline) founding a population will give rise to a population solely consisting of A2A2 descendants.

**Table S1.** *Icerya purchasi* microsatellite primer information for the markers used in this study. Throughout the supplement and main text, loci are referred to based on their primer label (PIcp#). Additional information on annealing temperatures can be found in the methods.

| primer label | primer code | F/R | sequence | Repeat | Dye | Multiplex | Size range |
|---|---|---|---|---|---|---|---|
| **PIcp21** | **I5YRX.81** | F | GAATGGAATGTGCGTTGTCT | $(TTA)_{11}$ | 6-FAM | 1 | 183-186 |
| | **I5YRX.81** | R | CAAGCTGATTTCCACTCTGTCTT | | | | |
| **PIcp39** | **IDTZN.140** | F | ATTCATCGAACCGCTTTCAA | $(ATA)_{13}$ | VIC | 1 | 165-176 |
| | **IDTZN.140** | R | AGCGCACTGTTAGCGTTGTA | | | | |
| **PIcp45** | **IMZ6I.153** | F | CGATGTATCTCGGTCGGAGT | $(TAT)_{15}$ | NED | 1 | 172-178 |
| | **IMZ6I.153** | R | CCCGTTTCTTTCAAGTCCAC | | | | |
| **PIcpP71** | **J0NJY.38** | F | CCCCCATTGATGCAAATAGT | $(TTA)_{12}$ | 6-FAM | 2 | 96 - 108 |
| | **J0NJY.38** | R | GTTGTTGCCTCGATGGAATG | | | | |
| **PIcp58** | **IUJEK.45** | F | CCGAATATAATTACGAAAGTAGCAAA | $(ATA)_{16}$ | VIC | 2 | 132 - 138 |
| | **IUJEK.45** | R | CGTTCGAAACATTCTGCGTA | | | | |
| **PIcp62** | **IWJ8O.147** | F | CAAAGCGAGATTTCCTGTCC | $(TTA)_{11}$ | NED | 2 | 169-176 |
| | **IWJ8O.147** | R | GGGATCCTCAAACGCAATAC | | | | |
| **PIcp75** | **JCKXY.97** | F | GCGCGTTAGTCAGATGAAGA | $(TTA)_{11}$ | PET | 2 | 154-160 |
| | **JCKXY.97** | R | CCTCCTCGCCCTTTCTTTAC | | | | |
| **PIcp101** | **JR7HX.82** | F | GTTTTTAGGACTGGCGGTGT | $(CTT)_{14}$ | 6-FAM | 4-1 | 98-107 |
| | **JR7HX.82** | R | AAGCATTGAGAGCAGCCAAC | | | | |
| **PIcp29** | **I6NB.264** | F | GGCACACTTTAATCCGAACG | $(ATA)_{19}$ | VIC | 4-1 | 125-128 |
| | **I6NB.264** | R | GGGCAAGGAACAGTCAAAGA | | | | |
| **PIcp68** | **IZ9OC.203** | F | CACCTGATTAGGCTATTGTTTATTT | $(ATT)_{13}$ | NED | 4-2 | 87-109 |
| | **IZ9OC.203** | R | CCCTAGAGAGATGCGAAGGA | | | | |
| **PIcp93** | **JNG87.33** | F | GAAGTATAAAGTGAAAGAAAGGAACTG | $(ATT)_{22}$ | PET | 4-2 | 114-120 |
| | **JNG87.33** | R | CGCGAGTGAGACTCAAGCTA | | | | |
| **PIcp37** | **IC578.156** | F | CGTGCGTTATTAAAGCCATTG | $(TAA)_{12}$ | VIC | 4-2 | 206-217 |
| | **IC578.156** | R | GTGCCGTTGGGCTTAGTTTA | | | | |

| PIcp21 | PIcp39 | PIcp45 | PIcpP71 | PIcp58 | PIcp62 | PIcp75 | PIcp101 | PIcp29 | PIcp37 | PIcp68 | PIcp93 |
|---|---|---|---|---|---|---|---|---|---|---|---|
| 144 | 162 | 137* | 90* | 99 | 149* | 132 | 72* | 102* | 206 | 104 | 115 |
| 171 | 165 | 143* | 93 | 108 | 152 | 147 | 95 | 105* | 209 | 107 | **121** |
| 180 | 168 | 146 | 96 | **131** | 170 | 150 | 98 | 114 | 212 | **110** | 124 |
| **183** | **171** | 167 | **99** | **134** | 173 | 153 | 101 | 120 | **215** | 122* | 127 |
| 186 | 174 | 170 | 102 | 137 | **176** | **159** | 104* | 123 | 221 | 125 | |
| | 177 | **173** | 105 | | 179 | 162 | **107** | **126** | 245* | 128 | |
| | | 176 | 108 | | | 165 | 122 | | 248 | | |
| | | 179 | 111 | | | 168 | 128 | | | | |
| | | 182 | | | | 171 | | | | | |
| **Total number off alleles** | | | | | | | | | | | |
| **3** | **5** | **7** | **7** | **4** | **5** | **6** | **7** | **4** | **5** | **4** | **2** |

**Table S2.** Allelic richness of loci examined in this study. Note that in spite of high levels of homozygosity, all loci included in analyses showed some variation globally. Numbers under each locus represent the size of the amplified microsatellite repeat fragment. Bolding and underlines denotes the most common allele for each locus. As mentioned in the main text, the two globally dominant genotypes differed at a single locus, PIcp58. Here the two most common alleles are noted. Asterisks denote private alleles in Australia.

| $F_{ST}$ | France | Turkey | Korea | Mexico | South Africa | Spain |
|---|---|---|---|---|---|---|
| California | 0.0328 | 0.0340 | 0.0674 | 0.0225 | 0.0595 | 0.0020 |
| France | NA | 0.0865 | 0.0920 | 0.0373 | 0.1118 | 0.04691 |
| Turkey | | NA | 0.0049 | 0.0671 | 0.0047 | 0.0577 |
| Korea | | | NA | 0.0627 | 0.001 | 0.0935 |
| Mexico | | | | NA | 0.0846 | 0.3547 |
| South Africa | | | | | NA | 0.0908 |

**Table S3.** Global pairwise $F_{ST}$ for each population included in the population structure analysis (i.e. those with more than one individual with genotypes at 8 or more of the 12 markers used here). Differentiation ranges from 0.001 (Korea-South Africa) to 0.11 (France-South Africa) based mainly on the presence of rare variants and shows no pattern of isolation by distance.

| Population code | Location | Country | Collection date | Host plant | Collector |
|---|---|---|---|---|---|
| CE | California | California, US | 2013 | *Citrus* | Mark Hoddle |
| AU | Canberra | Australia | 2012 | | Penny Gullan |
| BU | Bungonia | Australia | 30-1-2012 | *Acacia* | Mark Hoddle |
| ACP | Riverside | California, US | | | Mark Hoddle |
| CCS1 | University of CA Riverside | California, US | 12-11-10 | unknown | Mark Hoddle |
| CCS2 | Near CA campus residential Riverside | California, US | 31-12-2010 | *Pittosporum* | Mark Hoddle |
| R | Riverside | California, US | 27-12-2012 | *Citrus* | Mark Hoddle |
| SF | San Francisco, near Union Square | California, US | 2011 | *Pittosporum* | Laura Ross |
| TC | Trabuco Canyon | California, US | 2013 | *Orange/Grapefruit* | Mark Hoddle |
| FA | La Ciotat | France | 2011 | *Acacia* | |
| FP | La Ciotat | France | 2011 | *Pittosporum* | |
| FR | La Ciotat | France | 28-5-2011 | *Rosmarinus* | |
| MP | Montpellier | France | March 2012 | various | Laura Ross |
| KO | Jeju | Korea | | | Dong-Soon, Kim |
| MX | Acambaro, Guanajuato | Mexico | 30-1-2012 | *Citrus* | Mark Hoddle |
| PA | PARS, Faisalabad, Punjab | Pakistan | 8-4-2011<br>30-11-2011 | *Citrus* | Mark Hoddle |
| SA | Western cape region | South Africa | February-March 2013 | various | Mark Hoddle |
| SP | Valencia | Spain | 2011 | | |
| KAC | Cevlik, Hatey Province | Turkey | 20-9-2010 | *Citrus* | Mark Hoddle |
| KA | Kayhan, Adana area | Turkey | 16-9-2010 | *Citrus* | Mark Hoddle |
| VA | Vakif, Hatey province | Turkey | 20-9-2010 | *Citrus* | Mark Hoddle |
| CH | Chelsea flower show, London | UK | 2011 | unknown | |
| NH | Natural history museum London, wildlife garden | UK | September 2011 | *Gorse* | Laura Ross |

**Table S4.** Additional collecting information for samples, denoting country, location, and population code used in data files. When available, collection date and host plant on which the insects were found is also recorded; however, due to the collaborative nature of sample collecting from multiple countries, not all information was recorded for every population.

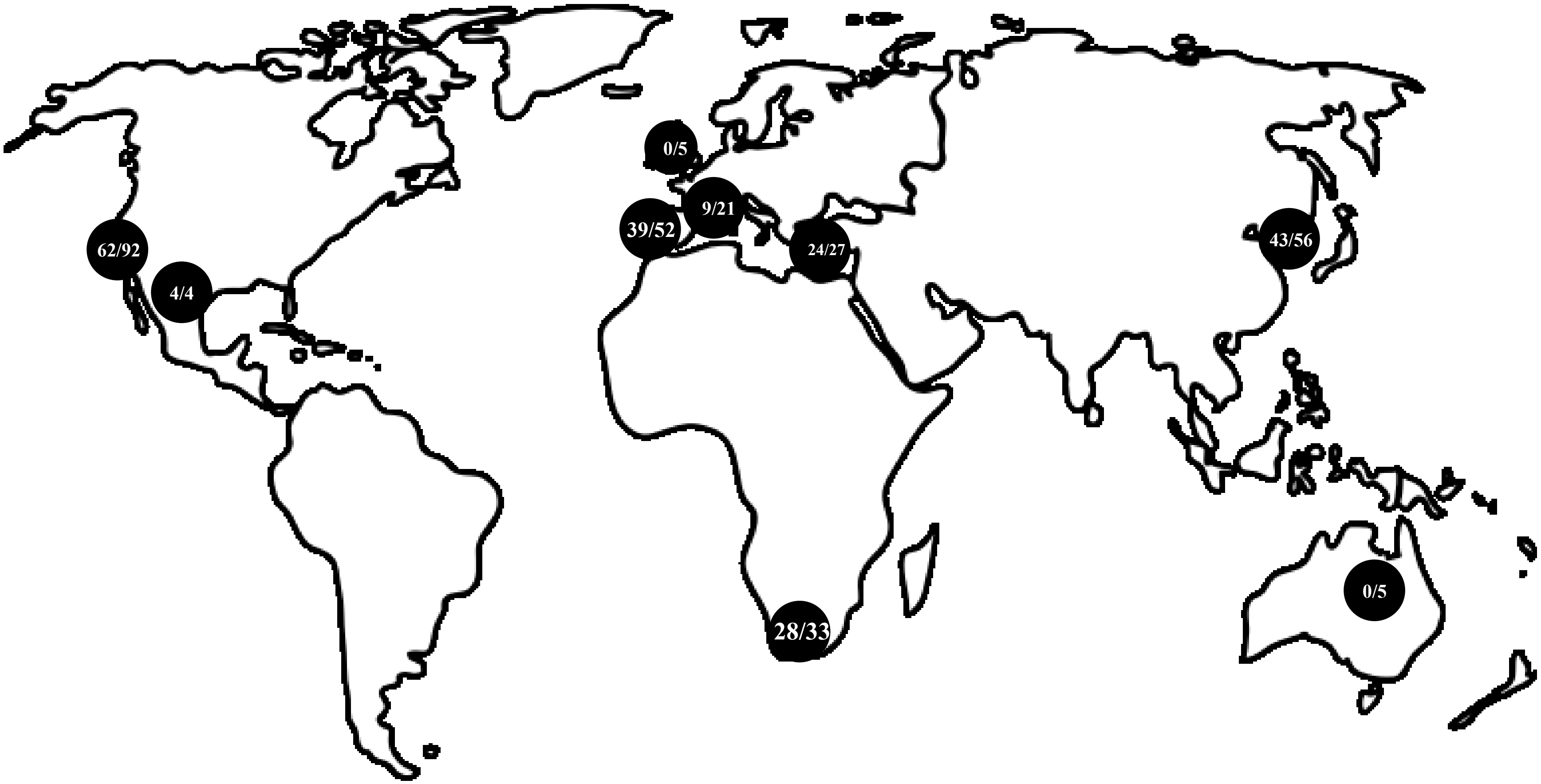


**Figure S1.** Map of all locations analyzed in initial collection of *Icerya purchasi*. Pairs of numbers follow the convention: first samples with complete genotypes used for $F_{ST}$, second samples with at least 8 amplified loci used in inbreeding analyses. For example, 28 South African samples were used to estimate $F_{ST}$, and 33 used to estimate inbreeding. In the case of Australia and the United Kingdom, none of the samples met cutoff criteria for $F_{ST}$ calculation.

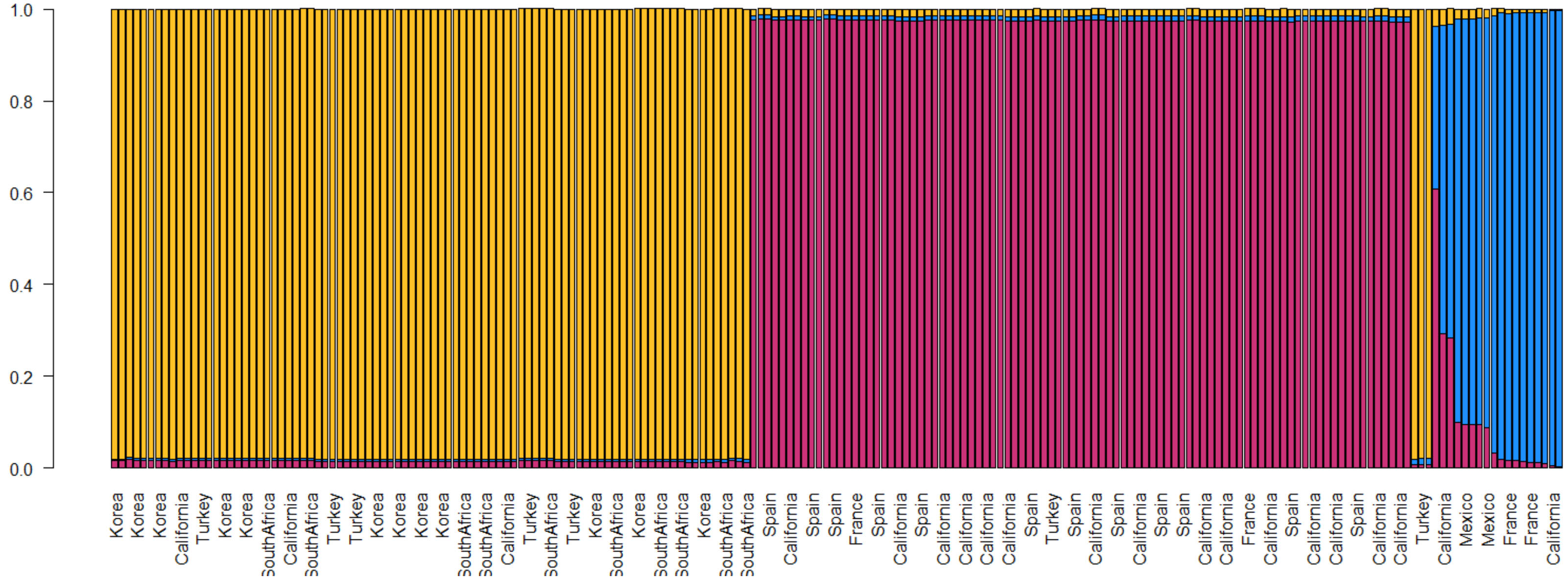


**Figure S2**. A representative structure plot for the most likely number of populations (k = 3). Individual bars represent individual insects, with colored proportions indicating the percent identity with one of the three inferred populations. Note that all individuals from one country often do not fall cleanly into a single cluster, likely owing to the high frequency of two nearly identical haplotypes that can be found across the world.
